# MAPK/ERK signaling in gliomas modulates interferon responses, T cell recruitment, microglia phenotype, and immune checkpoint blockade efficacy

**DOI:** 10.1101/2024.09.11.612571

**Authors:** Kwang-Soo Kim, Junyi Zhang, Víctor A. Arrieta, Crismita Dmello, Elena Grabis, Karl Habashy, Joseph Duffy, Junfei Zhao, Andrew Gould, Li Chen, Jian Hu, Irina Balyasnikova, Dhan Chand, Dan Levey, Peter Canoll, Wenting Zhao, Peter A. Sims, Raul Rabadan, Surya Pandey, Bin Zhang, Catalina Lee-Chang, Dieter Henrik Heiland, Adam M. Sonabend

**Affiliations:** Department of Neurological Surgery, Feinberg School of Medicine, Northwestern University, Chicago, IL, USA; Northwestern Medicine Malnati Brain Tumor Institute of the Lurie Comprehensive Cancer Center, Feinberg School of Medicine, Northwestern University, Chicago, IL, USA; Microenvironment and Immunology Research Laboratory, Medical Center - University of Freiburg, Freiburg, Germany; Department of Neurosurgery, Medical Center - University of Freiburg, Freiburg, Germany; Faculty of Medicine, University of Freiburg, Freiburg, Germany; Translational NeuroOncology Research Group, Medical Center - University of Freiburg, Freiburg, Germany; Department of Systems Biology, Columbia University, New York, NY, USA; Department of Biomedical Informatics, Columbia University, New York, NY, USA; Department of Cancer Biology, The University of Texas MD Anderson Cancer Center, Houston, TX, USA; The University of Texas MD Anderson Cancer Center UTHealth Graduate School of Biomedical Sciences, Houston, TX, USA; Agenus Inc., Lexington, MA, USA; Herbert Irving Comprehensive Cancer Center, Columbia University Irving Medical Center, New York, NY, USA; Department of Pathology and Cell Biology, Columbia University Irving Medical Center, New York, NY, USA; Department of Biochemistry & Molecular Biophysics, Columbia University, New York, NY, USA; Department of Hematology and Oncology, Northwestern University, Feinberg School of Medicine, Chicago, IL, USA

## Abstract

**Background:** Glioblastoma (GB) remains a formidable challenge in neuro-oncology, with immune checkpoint blockade (ICB) showing limited efficacy in unselected patients. We previously recently established that MAPK/ERK signaling is associated with overall survival following anti-PD-1 and anti-CTLA-4 treatment in recurrent GB. However, the causal relationship between MAPK/ERK signaling and susceptibility to ICB, as well as the mechanisms underlying this association, remain poorly understood.

**Method:** We conducted *in vivo* kinome-wide CRISPR/Cas9 screenings in murine gliomas to identify key regulators of susceptibility to anti-PD-1 and CD8^+^ T cell responses and performed survival studies to validate the most relevant genes. Additionally, paired single cell RNA-sequencing (scRNA-seq) with p-ERK staining, spatial transcriptomics on GB samples, and *ex-vivo* slice culture of a BRAF^V600E^ mutant GB tumor treated with BRAFi/MEKi were used to determine the causal relationship between MAPK signaling, tumor cell immunogenicity, and modulation of microglia phenotype.

**Results:** CRISPR/Cas9 screens identified the MAPK pathway, particularly the RAF-MEK-ERK pathway, as the most critical modulator of glioma susceptibility to CD8^+^ T cells, and anti-PD-1 across all kinases. Experimentally-induced ERK phosphorylation in gliomas enhanced survival with ICB treatment, led to durable anti-tumoral immunity upon re-challenge and memory T cell infiltration in long-term survivors. Elevated p-ERK in glioma cells correlated with increased interferon responses, antigen presentation and T cell infiltration in GB. Moreover, spatial transcriptomics and scRNA-seq analysis revealed the modulation of interferon responses by the MAPK/ERK pathway in BRAF^V600E^ human GB cells with ERK1/2 knockout and in slice cultures of human BRAF^V600E^ GB tissue. Notably, BRAFi/MEKi treatment disrupted the interaction between tumor cells and tumor-associated macrophages/microglia in slice cultures from BRAF^V600E^ mutant GB.

**Conclusion:** Our data indicate that the MAPK/ERK pathway is a critical regulator of GB cell susceptibility to anti-tumoral immunity, modulating interferon responses, and antigen-presentation in glioma cells, as well as tumor cell interaction with microglia. These findings not only elucidate the mechanistic underpinnings of immunotherapy resistance in GB but also highlight the MAPK/ERK pathway as a promising target for enhancing immunotherapeutic efficacy in this challenging malignancy.

## Introduction

Glioblastoma (GB), the most common and malignant primary brain tumors in adults, is characterized by pronounced genetic and molecular heterogeneity^1,2^. This diversity not only drives oncogene processes like cell proliferation but also leads to variations in the activation of key signaling pathways, such as MAPK and PI3K/AKT/mTOR^3,4^. The complexity of GB biology is further amplified by the numerous genetic and epigenetic alterations that can modulate these pathways, resulting in a spectrum of molecular profiles across tumors^5^. Moreover, the tumor microenvironment, plays a crucial role in GB progression, and in particular, tumor-associated macrophages and microglia (TAM) constitute up to 30-40% of the tumor volume^6^. These TAMs exhibit a phenotypic spectrum modulated by tumor cells, which varies across individual GB cases due to the underlying molecular heterogeneity^7^. This complex interplay between tumor cells and their microenvironment contributes significantly to the challenges in treating GB.

The molecular heterogeneity of GB significantly impacts clinical outcomes by contributing to the variability in treatment response^1,8–11^. In the context of immunotherapy, the variable tumor microenvironment likely contributes to inconsistent therapeutic outcomes^11^. Tailoring immunotherapy to account for the molecular diversity of GB could enhance efficacy in selected cases, even though traditional randomized controlled trials in unselected populations may not show overall survival benefits^12^. Anti-PD-1 (aPD-1) immunotherapy exemplifies this issue, as some cases demonstrate durable responses and prolonged survival, while multiple randomized controlled trials in unselected patients have failed to meet primary the efficacy endpoint^12–16^.

We reported an association between mutations that activate MAPK pathway (*PTPN11*/*BRAF*) and GB susceptibility to aPD-1 therapy^10^. Notably, even in the absence of these mutations, MAPK activation, determined by the phosphorylation of the ERK protein (p-ERK), predicted overall survival in GB patients undergoing aPD-1 and anti-CTLA-4 (aCTLA-4) therapy across several independent cohorts^9,17^. Tumors with elevated p-ERK^+^ tumor cell density exhibited a distinct microglial phenotype, characterized by the proximity of microglia to p-ERK^+^ tumor cells and the expression of MHC class II antigen presenting molecules^9^. However, the causal relationship between MAPK signaling, this microglial phenotype, and the observed susceptibility to immunotherapy remains unsolved. Moreover, the mechanism by which tumor cells with active MAPK signaling become more immunogenic or modulate adjacent TAM are unclear.

In this study, we leveraged large *in vivo* CRISPR knockout (KO) screens, mouse glioma models, *in vitro* experiments and reverse translation data derived from analyses of human tumors to investigate the impact of tumor cell-intrinsic p-ERK and MAPK signaling on the tumor cell immunogenicity, microglial phenotype and susceptibility to ICB. By elucidating the role of MAPK signaling in shaping tumor immunogenicity and the microenvironment, our findings provide a foundation for more precise and effective immunotherapy strategy tailored to the molecular profile of glioblastoma.

## Results

### *In vivo* kinome-wide CRISPR/Cas9 screens identify MAPK/ERK signaling as a regulator for susceptibility to CD8^+^ T cells and anti-PD-1 immunotherapy

To investigate whether MAPK/ERK signaling contributes to immune cell recognition of glioma cells, we conducted kinome-wide CRISPR/Cas9 screens. GL261 glioma cells were transduced with a lentiviral sgRNA library resulting in the knockout (KO) of a single kinase gene per tumor cell. These transduced cells were then implanted intracranially into mice (Figure 1A). By comparing the implanted glioma cells into wildtype and *Cd8* KO mice, we identified kinase KO clones that were selected in the presence of CD8^+^ T cells in the host^18^. Gene ontology (GO) analysis revealed the MAPK pathway as the most significantly enriched among all kinase pathways, in KO clones selected by CD8^+^ T cells in wildtype mice, compared to *Cd8* KO hosts (Figure 1B). Among MAPK-related genes, *Map2k2* (encoding Mek2) and *Araf*, both upstream of Erk phosphorylation, showed the highest enrichment (Figure 1C and D).

**Figure 1.**
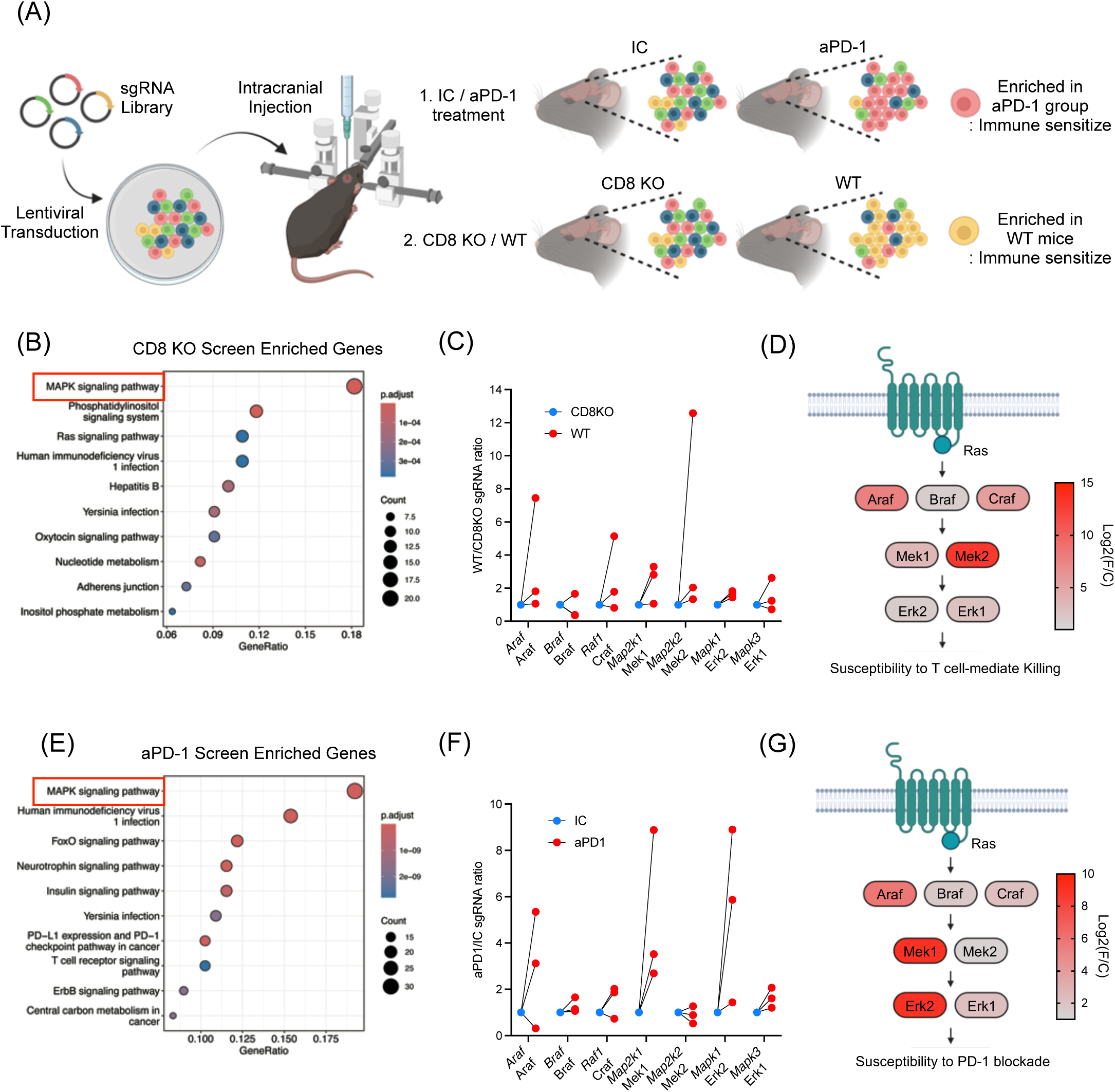
*In vivo* kinome-wide CRISPR/Cas9 screens identify MAPK/ERK signaling as a regulator for susceptibility to CD8^+^ T cells and anti-PD-1 immunotherapy. (A) Schematic illustration of *in vivo* CRISPR/Cas9 screening. SgRNA library transduced to GL261 mouse glioma cells and the cells were injected intracranially. Comparisons were 1) IC (n = 11) vs. aPD-1 (n = 12), 2) wildtype (n = 12) vs. *Cd8* KO mice (n = 20). (B) Gene enrichment analysis from the CRISPR/Cas9 screenings comparing wildtype and *Cd8* KO mice. (C) Fold changes of sgRNAs targeting MAPK/ERK pathway genes. Comparing wildtype and *Cd8* KO mice, 3 sgRNAs are depicted. (D) Detailed diagrams of the MAPK signaling cascade, indicating log fold changes in sgRNA enrichment. Log2(foldchange) was calculated based on the top performing sgRNA depleted in *Cd8* KO mice. (E) Gene enrichment analysis from CRISPR/Cas9 screening comparing mice received IC and aPD-1. (F) Fold changes of sgRNAs targeting MAPK/ERK pathway genes. Comparing IC and aPD-1 group, 3 sgRNAs are depicted. (G) Diagrams of the MAPK signaling cascade, indicating log fold changes in sgRNA enrichment. Log2(foldchange) was calculated based on the top performing sgRNA enriched in aPD1 treated group. For B and E, the gene list of the kinome was used as background (denominator) to calculate the gene ontology enrichment.

To access whether these kinases, identified as critical for glioma recognition by CD8^+^ T cells, also conferred susceptibility to ICB, we performed a similar CRISPR/Cas9 screen in which transduced glioma cells were implanted into wildtype mice treated with aPD-1 or with isotype control (IC) antibody. In this screen, the MAPK signaling pathway again emerged as the most enriched among all KO clones selected by aPD-1 treatment (Figure 1E). Within the MAPK pathway, genes such as *Araf*, *Raf1*, *Map2k1* (encoding Mek1), and *Mapk1* (encoding Erk2) showed the highest enrichment under aPD-1 treatment (Figure 1F and G), all involved in ERK activation. Consistent with previous reports, *Cd8* KO exhibited significantly reduced survival, while ICB treatment showed no survival difference in the GL261 mouse model (Supplementary Figure 1)^18,19^. Normalized sgRNA counts for each of these CRISPR screens are available in Supplementary Table 1.

### MAPK activation enhances glioma susceptibility to aPD-1 therapy and promote durable anti-tumoral immunity

To further investigate a causal link between MAPK signaling in glioma cells and anti-tumoral immunity, we overexpressed *Mek1* and *Mek2* in GL261 mouse glioma cells (GL261-Mek1/2), which led to increased Erk phosphorylation (GL261-Mek1/2 p-Erk^high^; Figure 2A and B). Intracranial implantation of GL261-Mek1/2 p-Erk^high^ cells, resulted in tumors with higher CD8^+^ T cell infiltration compared to controls (Figure 2C), consistent with our earlier observation that KO clones for *Mek1* and *Mek2*, were selected in the presence of CD8^+^ T cells (Figure 1C and D). Tumors derived from GL261-Mek1/2 p-Erk^high^ cells exhibited marked susceptibility to aPD-1 immunotherapy, with over 60% of the mice achieving long-term survival (P = 0.001, Figure 2D). This effect was CD8^+^ T cell-dependent, as *Cd8* KO hosts bearing GL261-Mek1/2 p-Erk^high^ tumors showed no survival benefit from aPD-1 treatment (Figure 2E).

**Figure 2.**
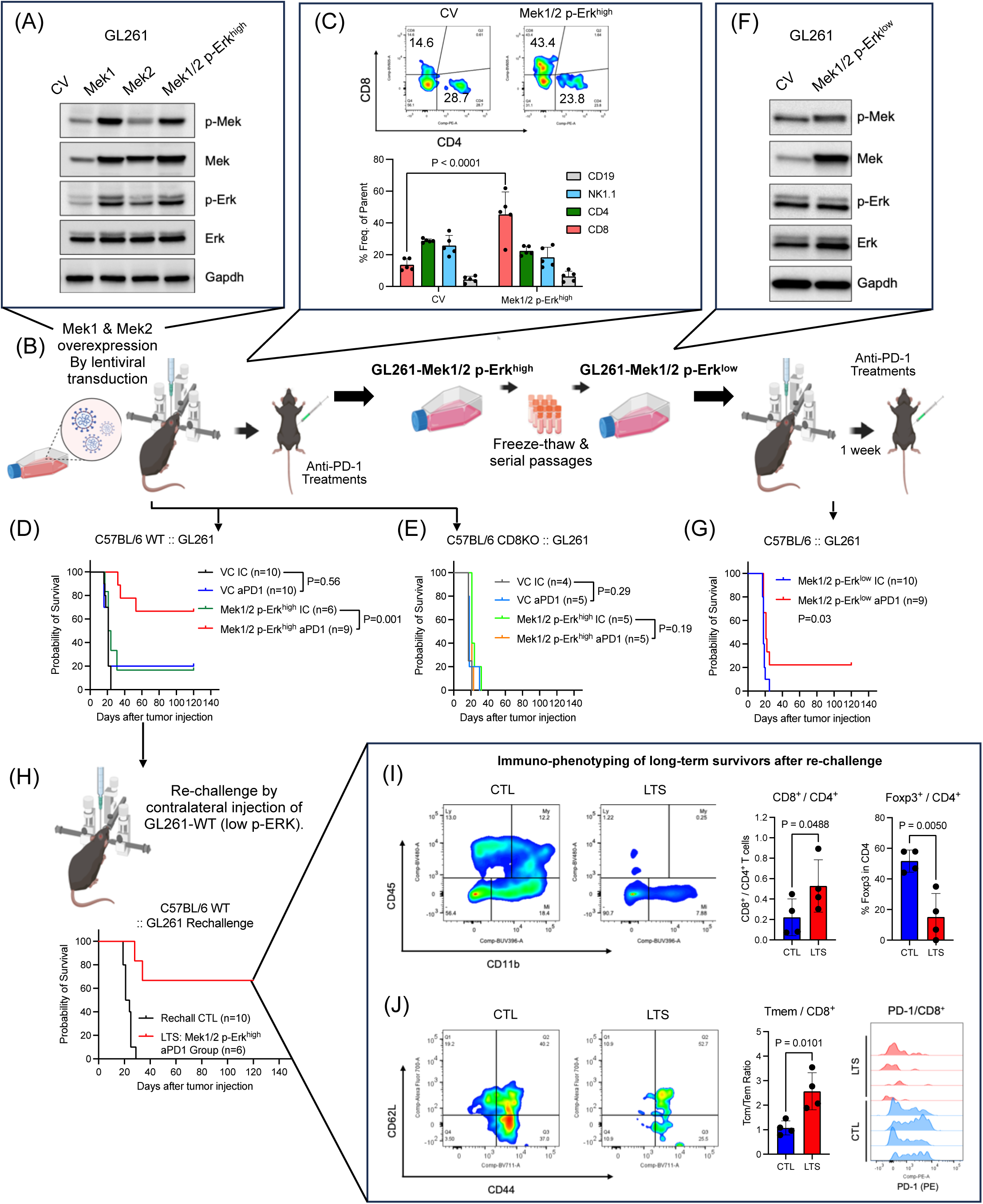
MAPK/ERK activation in glioma cell contributes to susceptibility and durable anti-tumoral immunity following anti-PD-1 therapy. (A and B) Schematic illustration of cell line generation and *in vivo* survival studies for GL261 Mek1/2 overexpression stable cell line was established by lentiviral transduction and confirmed by Western blot to confer Erk1/2 phosphorylation. For treatment with ICB antibodies, treatment started 1 week after tumor implantation and delivered intravenously 4 times every 3 or 4 days. (C) Flow cytometry based immune cell infiltration analysis. After 20 days of tumor cell infusion, tumor infiltrating lymphocytes were analyzed comparing control GL261 tumor (n = 5) and GL261-Mek1/2 p-Erk^high^ tumor (n = 5). (D) Kaplan-Meier survival curve of GL261-CV (IC n = 10, aPD-1 n = 10) and GL261-Mek1/2 p-Erk^high^ (IC, n = 6 and aPD-1, n = 9) treated with IC or aPD-1 antibody, conducted in wildtype C57BL/6 mice. (E) Kaplan-Meier survival curve of control GL261-CV (IC, n = 4 and aPD-1, n = 5) and GL261-Mek1/2 p-Erk^high^ (IC, n = 5 and aPD-1, n = 5) treated with IC or aPD-1 antibody, conducted in *Cd8* KO C57BL/6 mice. (F) Western blot analysis of Erk and Mek phosphorylations and expressions after several passages and freeze/thaw cycles. The cell line was referred as GL261-Mek1/2 p-Erk^low^. (G) Kaplan-Meier survival curve of control GL261 tumor and GL261-Mek1/2 p-Erk^low^ treated with IC (n = 9) or aPD-1 (n = 10), conducted in wildtype C57BL/6 mice. (H) The mice cohort from GL261-Mek1/2 p-Erk^high^, the long-term survivors received contralateral injection of wildtype GL261 cell infusion on day 120 post tumor cell injection (control, n = 10 and long-term survivors, n = 6). (I) Immune landscape after 120 days of the rechallenge. Comparing with brand-new tumor (CTL, n = 4), brains from long-term survivors (LTS, n = 4) were analyzed for immune phenotyping. Flow cytometry data showing percoll-enriched cells (left). Bar graph showing CD8^+^ and CD4^+^ T cell ratio (middle), Foxp3^+^ cells in CD4^+^ T cell population (right). (I) Flow cytometry data showing expression of CD44 and CD62L in CD8^+^ T cells (left). Bar graph showing central memory T cell (CD44^+^CD62L^+^) to effector memory T cell (CD44^+^CD62L^-^) ratio (middle). Flow cytometry data showing PD-1 expression in CD8^+^ T cells. The P values for survival studies were generated from low-rank test. Statistical analysis for group comparison was done using unpaired two-tailed T test.

Interestingly, after multiple freeze-thaw cycles and several passages in culture, GL261-Mek1/2 cells maintained overexpression of *Mek1* and *Mek2* but lost ERK phosphorylation (Figure 2F). When implanted intracranially, these cells with diminished p-Erk levels (GL261-Mek1/2 p-Erk^low^), showed reduced susceptibility to aPD-1 therapy, with no increase in median survival compared to isotype control-treated mice, and only 22.2% of mice achieving long-term survival (Figure 2G), further supporting the specific role of MAPK activation in immunotherapy response.

To further characterize the durability of anti-tumoral immunity, we re-challenged long-term survivors with parental GL261 cells implanted on the contralateral side of the brain. Notably, four out of six mice rejected the secondary tumors without any additional therapy (Figure 2H), suggesting that the anti-tumoral immune response was related to MAPK activation and directed against endogenous tumor antigens, rather than introduced transgenes. Immune phenotyping of long-term survivors revealed increased CD8/CD4 T cell ratio (P = 0.0488), decreased intra- tumoral FoxP3^+^ regulatory T cells (Tregs) (P = 0.0050, Figure 2I), a higher ratio of central memory to effector memory CD8^+^ T cells (P = 0.0101) and reduced PD-1 expression on infiltrating CD8^+^ T cells (Figure 2J).

We further validated these findings in a transgenic-derived murine glioma model QPP7, known for its susceptibility to immunotherapy^20^. These cells exhibited elevated p-Erk levels and targeted KO of Erk1 (QPP7-Erk1 KO) and Erk2 (QPP7-Erk2 KO) reduced the efficacy of aPD-1 therapy compared to non-target controls (Supplementary Figure 2). Median survival difference between IC-treated and aPD-1-treated groups was 7 days for QPP7-Erk1 KO (aPD-1: 64 days; IC: 57 days) and 13 days for QPP7-Erk2 KO (aPD-1: 57 days and IC: 44 days) (Supplementary Figure 2). In contrast, mice bearing QPP7-NTC tumors and treated with aPD-1 did not reach the median survival endpoint, further supporting a potential role of MAPK/ERK signaling in immunotherapy response (Supplementary Figure 2). Together, these data demonstrate that MAPK activation in glioma cells enhances susceptibility to aPD-1 therapy and promotes durable, tumor-specific immunity, primarily mediated by CD8^+^ T cells.

### MAPK/ERK activation improves the efficacy of Fc-enhanced anti-CTLA-4 immune checkpoint blockade in murine gliomas

Our CRISPR/Cas9 screens suggested that MAPK activation broadly impacts anti-tumoral immunity in gliomas. To determine whether this effect extends beyond aPD-1 therapy, we examined the impact of MAPK activation on the efficacy of aCTLA-4 immunotherapy. While conventional aCTLA-4 therapy has not resulted in significant survival benefits for unselected GB patients^16^, our previous studies suggest that elevated tumor p-ERK levels could be a key determinant of therapeutic response and prolonged survival in GB patients treated with a combination of aPD-1 and aCTLA-4 blockade therapy^19,21^.

To investigate whether MAPK activation also contributes to glioma susceptibility to aCTLA4 therapy, we used a mouse surrogate of botensilimab, a next generation Fc-enhanced aCTLA-4 antibody designed to bind with high affinity to activating Fcγ receptors expressed by host immune cells and extend anti-tumor immunity against ‘cold’ tumors^22–24^. We recently reported that this antibody is superior to conventional aCTLA-4 in glioma models^19^, and is currently being evaluated in an ongoing clinical trial in patients with GB (NCT05864534). In the GL261 mouse glioma model, resistant to both conventional and Fc-enhanced aCTLA-4^19^, activation of the MAPK signaling led to improved efficacy of Fc-enhanced aCTLA-4 therapy. Mice bearing GL261 Mek1/2 p-ERK^high^ tumors treated with the Fc-enhanced aCTLA-4 antibody achieved a 100% survival rate, which was significantly better than that observed in vector control tumors treated with the same antibody (P = 0.0291), and to Mek1/2 p-ERK^high^ tumors treated with isotype control antibody (P = 0.0012) (Figure 3). These findings highlight MAPK/ERK activation as a key enhancer of glioma susceptibility to both aPD-1 and Fc-enhanced aCTLA-4 therapies, suggesting a broader role in modulating tumor responsiveness to ICB therapy and its potential as a targeted therapeutic strategy for glioblastoma.

**Figure 3.**
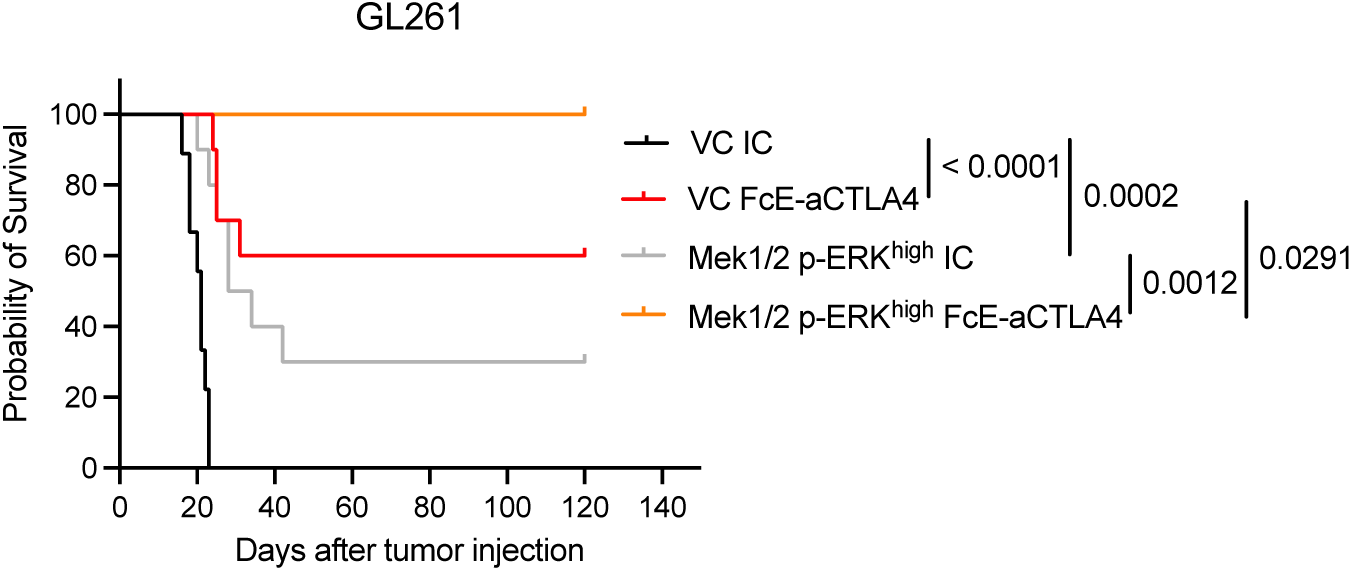
Activation of MAPK/ERK pathway is associated with the efficacy of Fc- enhanced anti-CTLA-4 therapy. Kaplan-Meier survival curve of GL261-CV (IC, n = 9 and Fc- enhanced aCTLA-4, n = 10) and GL261-Mek1/2 p-Erk^high^ (IC, n = 10 and Fc-enhanced aCTLA- 4, n = 10) treated with IC or Fc-enhanced aCTLA-4 antibody. For treatment with ICB antibodies, treatment started 1 week after tumor implantation and delivered intravenously 4 times every 3 or 4 days. The P values were generated from low-rank test.

### MAPK/ERK pathway is associated with glioblastoma cell response to Interferons

To elucidate the relationship between cell-intrinsic MAPK/ERK signaling and immune-related tumor phenotypes in GB, we conducted a continuous paired analysis of scRNA-seq data from GB patients with quantified p-ERK^+^ cell density (Figure 4A). This analysis revealed correlations between p-ERK^+^ cell density and gene expression of signaling pathways known to interact with MAPK/ERK, including receptor tyrosine kinase (RTK), WNT, AMP-activated protein kinase, and HIF-1⍺ pathways (Supplementary Figure 3)^3,25–27^. Notably, both continuous and dichotomous analyses identified significant correlations between p-ERK^+^ cell density and gene ontology (GO) themes associated with interferon (IFN) responses (Figure 4A and Supplementary Figure 3).

**Figure 4.**
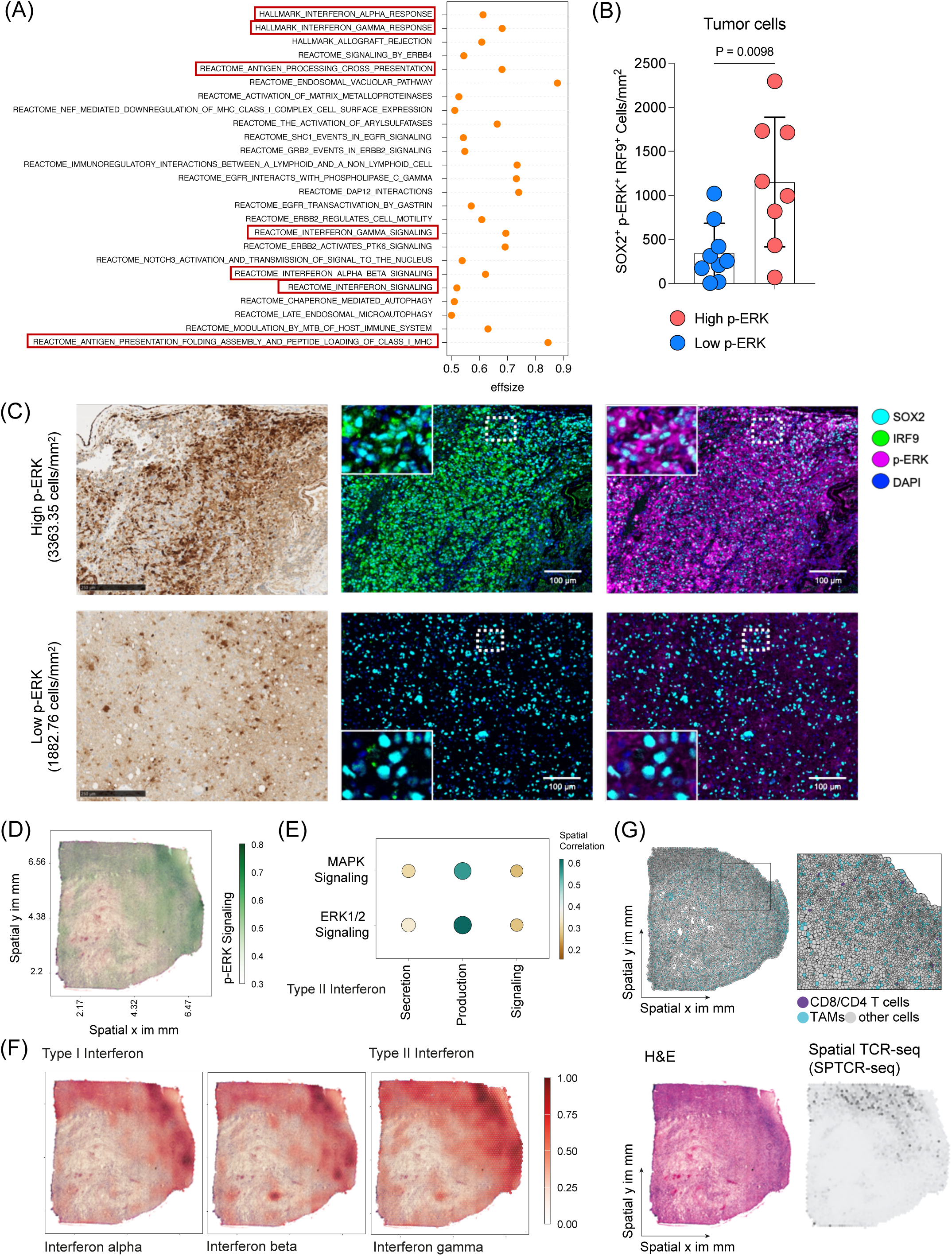
MAPK/ERK pathway is associated with interferon signaling in GB. (A) Differentially expressed gene signatures of tumor cells from single cell RNA sequencing comparing high and low p-ERK GB tumors. (B) Quantification analysis of immunofluorescence analysis representing SOX2^+^p-ERK^+^IRF9^+^ cell density in high p-ERK (n = 8) and low p-ERK (n = 9) tumors. Unpaired two-tailed T test was used, and P-value is depicted. (C) Representative immunohistochemistry (IHC) and immunofluorescence (IF) images of high p-ERK and low p- ERK GB patients depicting p-ERK and IRF9 expression. Scale bar = 250 µm for IHC and 100 µm for IF. (D) Example image of spatial multi-omic analysis of GB tumor representing p-ERK signaling activity. (E) Spatial correlation analysis of GB tumors showing correlation between MAPK and ERK1/2 signaling and type II IFN signatures. (F) Representative images of spatial muli-omic analysis demonstrate Type I and II interferon responses in GB tumor. (G) Single-cell composition after deconvolution representing the location of T cells and TAMs (upper). H&E image demonstrates the histology of GB tumor, and the image of spatial T cell receptor sequencing shows infiltrating T cells in GB tumor (low).

To validate the association of p-ERK with IFN responses at the cellular level, we employed multiplex immunofluorescence analysis. We compared the GB microenvironments between tumors with high (>3000 cells/mm^2^) and low p-ERK density, staining for SOX2 to identify GB cells, p-ERK to indicate MAPK/ERK activity, and IRF9, a marker for IFN responses^28^ (Figure 4B and C). IRF9 expression primarily localized to SOX2^+^ tumor cells (Supplementary Figure 4). Notably, aligning with the transcriptomic results, SOX2^+^IRF9^+^ cells/mm^2^ correlated with SOX2^+^p-ERK^+^ cells across tumors (R = 0.62, P = 0.0104). Tumors with high p-ERK exhibited a significantly higher density of SOX2^+^p-ERK^+^IRF9^+^ tumor cells compared to low p-ERK tumors (P = 0.0098, Figure 4B and C).

To investigate the spatial relationship between ERK signaling, IFN responses, and T cell abundance, we analyzed spatially resolved multi-omic data, including gene expression and T cell receptor sequencing (SPTCR-seq), across 12 primary GB samples^29^. Spatial analysis revealed a correlation between the MAPK/ERK signaling pathway and type II IFN secretion, production, signaling (Figure 4D and E). Both type I and II IFN signatures were co-localized with regions of activated p-ERK signaling (Figure 4F). Consistent with our *Cd8* KO CRISPR/Cas9 screen (Figure 1B) and *in vivo* immunophenotype analysis (Figure 2C), p-ERK signaling was closely associated with T cell abundance, as determined by single-cell deconvolution and SPTCR-seq (Figure 4G). Further, analysis of public RNA-seq data from patients who received either adjuvant or neoadjuvant PD-1 blockade^14^ suggested that high ERK signaling and corresponding type II IFN signaling were associated with improved survival (Supplementary Figure 5).

These findings collectively demonstrate a strong association between MAPK/ERK activation, interferon responses, and T cell infiltration in GB, providing mechanistic insights into the enhanced immunotherapy responsiveness observed in tumors with high MAPK/ERK activity.

### MAPK/ERK pathway drives interferon responses in glioma

To investigate the causal relationship between MAPK/ERK signaling and the observed tumor cell phenotypes across tumors, RNA-seq data from AM38, a human GB cell line with BRAF^V600E^ mutation, was analyzed following CRISPR-based single-gene KO of ERK1 and ERK2 (Supplementary Figure 6A). Given the strong association between the ERK pathway and cell growth^30,31^, ERK-deficient AM38 cells exhibited slower growth compared to non-target control (NTC) AM38 cells (Supplementary Figure 6B).

Differential gene expression analysis comparing AM38-NTC with AM38-ERK1 KO and AM38-ERK2 KO revealed GO themes modulated by MAPK/ERK signaling, including antigen processing MHC class I, and IFN responses (Figure 5A and B). Gene set enrichment analysis (GSEA) showed significant downregulation of type I and type II IFN responses genes in ERK1/2 KO cells, implicating ERK1/2 signaling in maintaining these immune-related tumor phenotypes (Figure 5B).

**Figure 5.**
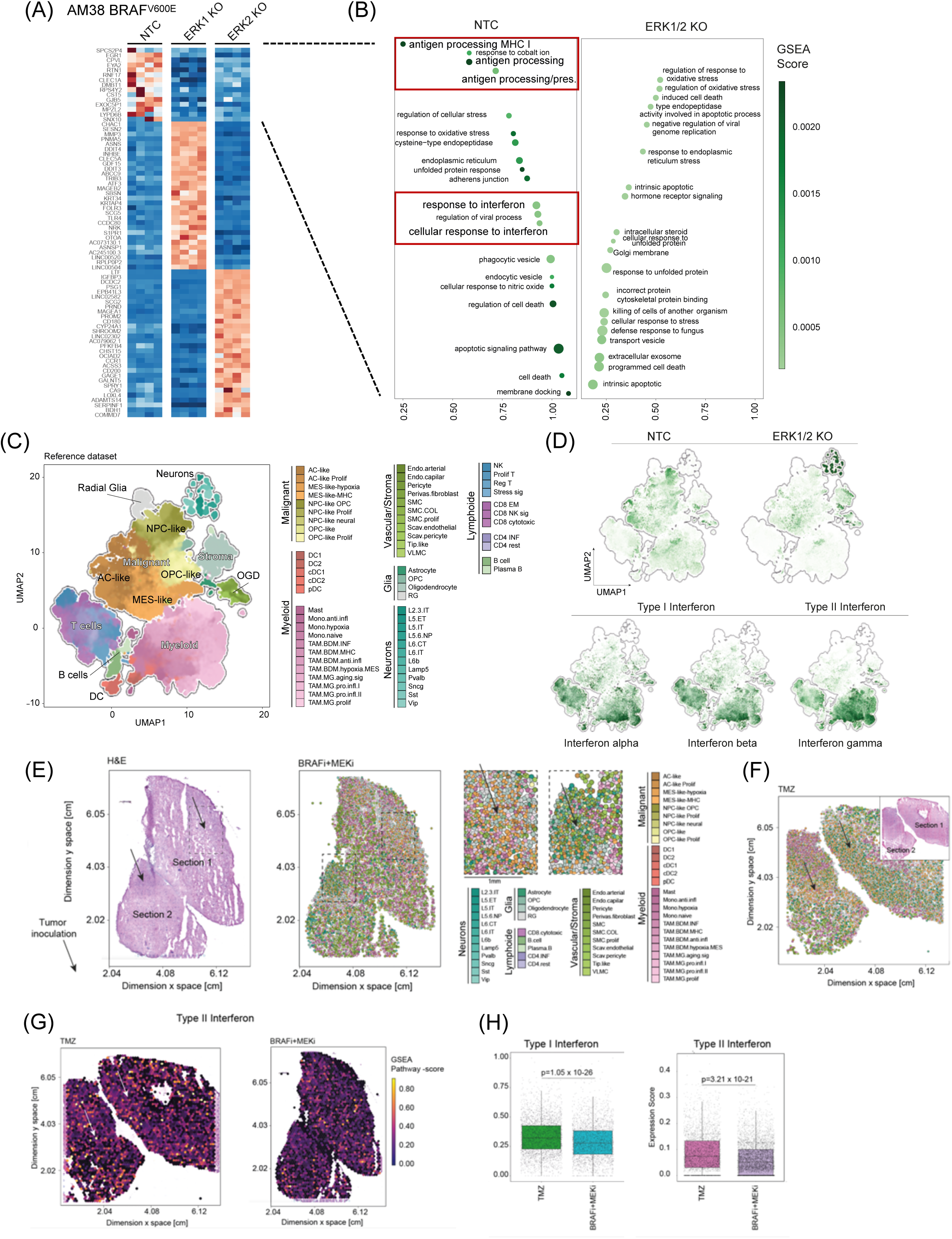
MAPK/ERK pathway confers interferon responses in glioma. (A) Bulk RNA- seq analysis of human GB cell line AM38-NTC and AM38-ERK KO. Heatmap displaying differential gene expression. NTC vs ERK1/2 KO is focused for further analysis. (B) Gene set enrichment analysis comparing AM38-NTC and AM38-ERK1/2 KO. (C) Uniform manifold approximation and projection dimensional reduction of the GBmap reference dataset. Each color indicates the different cell types. (D) Gene signatures from AM38-NTC and ERK1/2 KO are projected on GBmap reference dataset (top)^33^. Signatures of interferon alpha, beta, and gamma in the GBmap reference dataset (bottom). (E) Spatial transcriptomic and scRNA-seq analysis of slice culture of BRAF^V600E^ GB tumor treated with BRAFi/MEKi. The H&E image demonstrates the histology of slice culture (left). Each color indicates the different cell types (right). (F) Spatial transcriptomic and single cell RNA sequencing analysis of slice culture GB samples treated with temozolomide. (G) Spatial images of Type II interferon signature in slice culture samples treated with temozolomide (left) and BRAFi/MEKi (right). (H) Differential analysis of interferon type I and II signatures comparing temozolomide and BRAFi/MEKi treated slice culture.

Based on these findings, the potential for MAPK/ERK activation to enhance glioma cell responsiveness to type I IFN was explored. Functional analysis of IFN-α responsiveness in AM38 cells revealed that ERK1/2 KO significantly reduced the upregulation of type I IFN response genes (*IRF7*, *IRF9*, and *ISG15)*^32^ following IFN-α exposure (Supplementary Figure 7A). Given our previous observation that p-ERK in GB cells is associated with a distinct microenvironment phenotype^9^, the effect of ERK1/2 KO on cytokine secretion in response to IFN-α was further examined. While AM38 secrete CCL3/4 and GM-CSF following IFN-α exposure, this secretion was attenuated in AM38 ERK1 KO and ERK2 KO cells (Supplementary Figure 7B).

Analysis of a large public GB scRNA-seq dataset (GBmap)^33^ revealed differential expression of gene signatures derived from AM38-NTC and AM38-ERK1/2 KO across various cell populations (Figure 5C). The AM38-NTC signature (associated with elevated p-ERK, Supplementary Figure 6A) showed enhanced expression within tumor cells, particularly astrocyte (AC)-like and neural progenitor cell (NPC)-like populations. Conversely, the AM38-ERK1/2 KO signature was predominantly enriched in the mesenchymal (MES)-like cell population, whereas IFN molecules, including IFN-α, IFN-β, and IFN-γ, are primarily expressed by T cells and myeloid cells (Figure 5D).

To explore the causal relationship between MAPK/ERK signaling and these phenotypes in human GB tissues, we utilized a novel autologous human neocortical slice model from a BRAF^V600E^ mutated GB patient (Supplementary Figure 8A)^34^. BRAF^V600E^-mutated tumor cells were implanted into cortex slices and treated with either temozolomide (TMZ) or BRAF/MEK inhibitors (BRAFi/MEKi). The optimal BRAFi/MEKi combination was determined based on maximal inhibition of tumor growth (Supplementary Figure 8B). Spatially resolved transcriptomic and scRNA-seq analyses of the slices conducted 7 days after treatment (Figure 5F and G), revealed downregulation of IFN signatures in BRAFi/MEKi-treated samples compared to TMZ treatment (Figure 5H).

### MAPK/ERK signaling modulates antigen-presenting molecules in glioma cells

Earlier analysis of scRNA-seq data from GB specimens with high and low p-ERK levels highlighted GO signatures related to antigen presentation (Figure 4A). Additionally, RNA-seq data from AM38 cells with ERK1/2 KO indicated that MAPK/ERK signaling modulates transcriptional signatures associated with antigen processing and presentation (Figure 5B). To determine whether this p-ERK-associated phenotype is present in human GB, multiplex immunofluorescence was used to assess the expression of antigen-presenting molecules. While HLA-DR was broadly expressed across cell types (Supplementary Figure 9), the density of SOX2^+^HLA-DR^+^p-ERK^+^ cells was significantly higher compared to SOX2^+^HLA-DR^+^p-ERK^-^ cells (P = 0.0011, Figure 6A and B).

**Figure 6.**
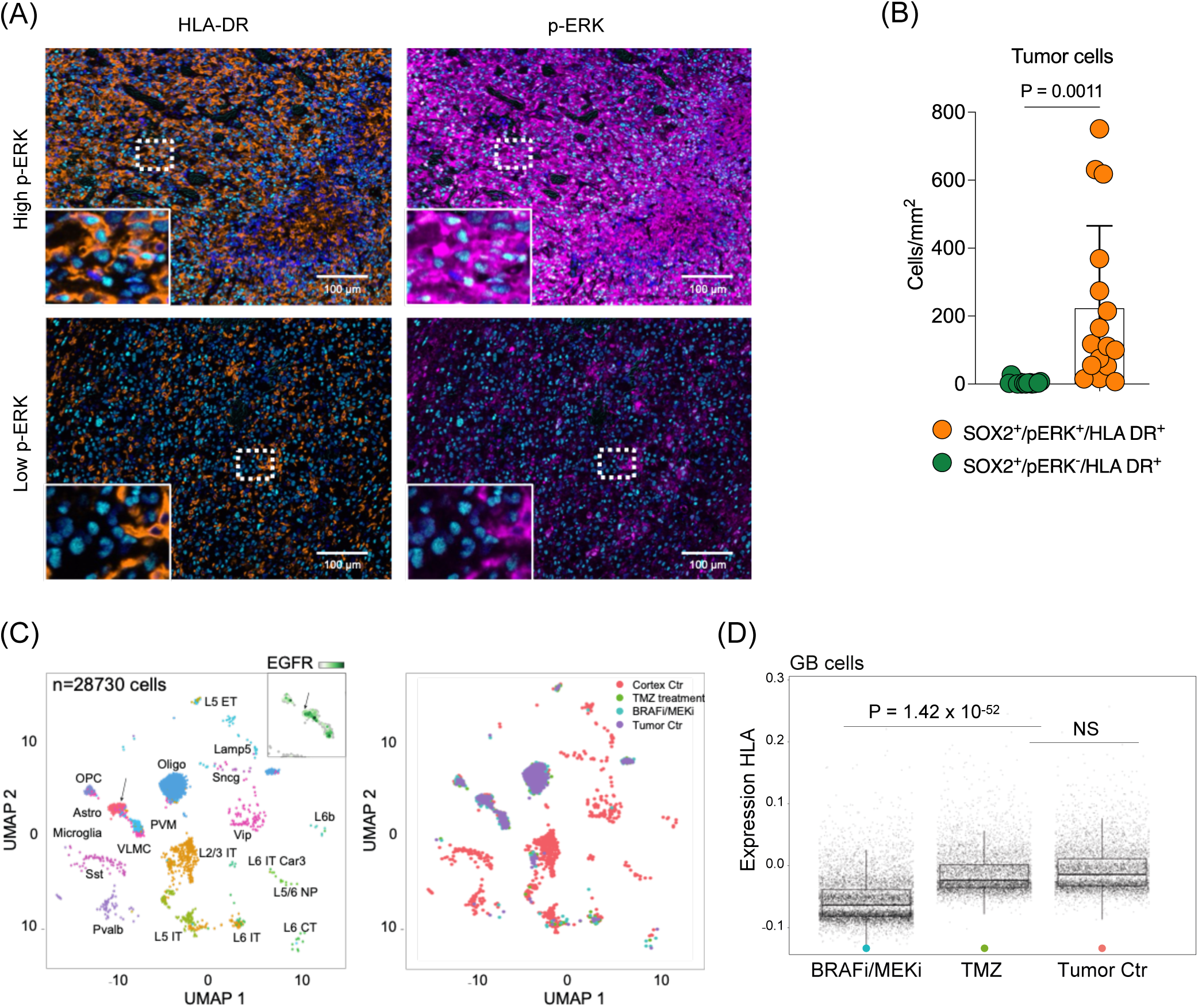
MAPK/ERK pathway modulates antigen presenting process in glioma cells. (A) Multiplex immunofluorescence analysis of human GB tumor. Representative images of high p- ERK tumor (top) and low p-ERK tumor (bottom). Scale bar = 100 µm. (B) Quantified phenotypic analysis of multiplex immunofluorescence across 16 GB patient samples. Bar graph showing SOX2^+^HLA-DR^+^ cells comparing p-ERK^-^ and p-ERK^+^ cells. (C) Single cell analysis from slice cultivated human GB tumor treated with temozolomide (left) and BRAFi/MEKi (right). (D) Differential expression of human leukocyte antigen (HLA) in slice cultivated human GB sample without treatment or temozolomide and BRAFi/MEKi treated (P = 1.42 x 10-52).

The causal relationship between MAPK/ERK activation and antigen presenting machinery expression in GB was further confirmed using the BRAF^V600E^ GB slice culture model. Here, scRNA-seq analysis revealed that BRAFi/MEKi treatment led to downregulation of HLA molecules in glioma cells, compared to TMZ treatment or no treatment (P = 1.42 x 10^-52^, Figure 6C and D). Moreover, *in vitro* studies using AM38 cells showed that ERK2 KO specifically led to downregulation of MHC II-related gene transcripts (Supplementary Figure 10A) and corresponding protein levels (Supplementary Figure 10B). Similar findings were observed in the murine glioma line QPP7, which exhibited elevated p-Erk, and susceptibility to aPD-1 therapy. Both QPP7-Erk1 KO and QPP7-Erk2 KO showed downregulation these genes (Supplementary Figure 10C and D).

These findings collectively demonstrate that MAPK/ERK signaling plays a crucial role in modulating the expression of antigen presenting molecules in glioma cells, providing a mechanistic link between MAPK activation and enhanced tumor cell immunogenicity.

### MAPK signaling in tumor cells drives activation of tumor-associated microglia in GB

Building on previous observations that high p-ERK gliomas exhibit a distinct microglia phenotype^9^, the causal contribution of MAPK/ERK signaling to GB cell-microenvironment interactions was further investigated using the slice culture model of BRAF^V600E^ GB. Graph-based cell proximity and communication analysis demonstrated that BRAFi/MEKi significantly reduced the communication between tumor cells and microglia. This effect was less pronounced for bone marrow-derived myeloid cells, indicating that MAPK signaling preferentialy contributes to cell-to-cell communication between tumor cells and tumor-infiltrating microglia (Figure 7A). Additionally, BRAFi/MEKi treatment modulated the microglial and TAM phenotypes. Notably, MAPK inhibition led to an increase in anti-inflammatory monocytes, and a decrease in proliferative microglia phenotype (Figure 7B). These findings demonstrate that MAPK /ERK signaling in GB cells plays a crucial role in shaping the tumor microenvironment, particularly in modulating microglia and TAM phenotypes and their interactions with tumor cells.

**Figure 7.**
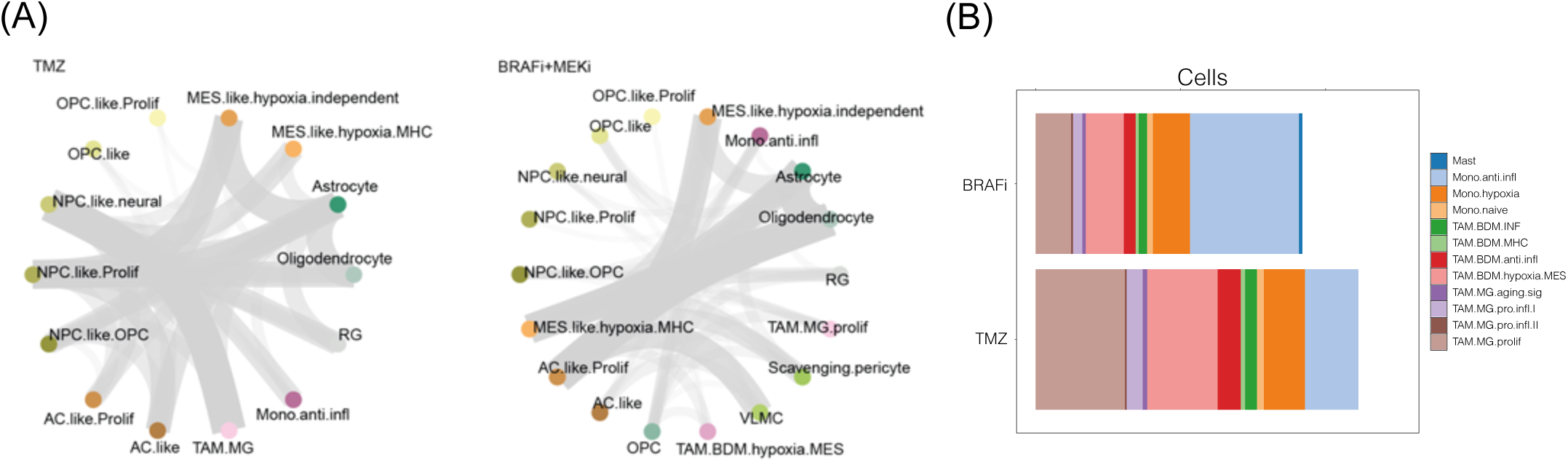
MAPK activation in GB tumor is associated with interaction between tumor cell and microglia. (A) Cell-to-cell communication analysis of slice cultivated human GB samples treated with temozolomide. (B) Phenotypic analysis of slice cultivated human GB samples treated with BRAFi/MEKi.

## Material and Method

### Orthotopic mouse glioma model

C57BL/6 mice were purchased from Charles River Laboratories and housed in a pathogen-free animal facility at Northwestern University Center for Comparative Medicine. All mouse work performed in this study were approved by the Institutional Animal Care and Use Committee (IACUC) under protocol number IS00014080. For tumor cell implantation, 6 to 8-week-old mice, were anesthetized with ketamine (100 mg/kg) and xylazine (10 mg/kg), and 100,000 glioma cells in 2.5 µl DPBS were injected intracranially at a specific brain coordinate after disinfecting and making a small cranial opening at 3 mm lateral and 2 mm caudal relative to the bregma using stereotaxic device (Harvard Apparatus). Post-surgery, animals were treated with isotype control or ICB antibodies and monitored until endpoints defined by the IACUC were met, including significant weight loss or severe neurological symptoms.

### Cell culture

The mouse glioma cell lines GL261 was obtained from the National Institutes of Health and cultured in Dulbecco’s modified Eagle’s medium supplemented with 10% fetal bovine serum and 1% penicillin/streptomycin. QPP7 cells^20,35^, kindly provided by Dr. Amy B. Heimberger from Northwestern University, were maintained in Dulbecco’s modified Eagle’s medium/F-12 (Gibco) supplemented with B-27 (Gibco), recombinant epidermal growth factor (EGF, 20 ng/ml, Peprotech), and recombinant basic fibroblast growth factor (bFGF, 20 ng/ml, Peprotech). All the cells were maintained in 5% CO2 incubators at 37℃ and tested for mycoplasma and confirmed negative before intracranial injection. Patient-derived GB cells after single cell suspension procedure were seeded in T-75 culture flasks and preserved in the cell culture incubator at 37 ℃ with 5% CO2 for recovery and following experiments. GM was used as cell culture medium for patient-derived GB cells in this study.

### Kinome-wide CRISPR Cas9 Screening

Both CD8KO and anti-PD-1 kinome-wide CRISPR screens were analyzed using the Model-based Analysis of Genome-wide CRISPR-Cas9 Knockout (MAGeCK) computational tool.^36^ Briefly, MAGeCK count command was used to map FASTq files to the sgRNAs in the Brie library. Counts were normalized to internal non-target sgRNA controls and gene essentiality scores (Beta-scores) was determined for each group relative to day 0 baseline controls using the MAGeCK maximum likelihood estimation (MLE) method.^37^ Kyoto Encyclopedia of Genes and Genomes (KEGG) Pathway enrichment analysis was performed on significant genes (FDR < 0.05) in R (v4.2.3)^38^ using clusterProfiler (v4.12.0)^39^ package and plots were generated using enrichPlot (v1.24.0).^40^

### Immunophenotype analysis

Tumor-bearing brans were processed immune profiling on 20 days post intra-cranial tumor injection after 2 treatments. Following preparation of Percoll gradient enriched single-cell suspension using a 70 µm cell strainer, cells underwent Fc blocking (TrueStain FcX, Biolegend) for 10 min for further staining process. Cells then underwent surface staining with primary antibodies and live/dead staining with Fixable Viability Dye eFluor 780 (eBioscience). After fixation and permeabilization (Foxp3/Transcription Factor Staining Buffer Set, eBioscience), intracellular staining was performed. Antibodies used in this study were listed in Supplementary Table 1. Flow cytometry data was acquired by the BD Symphony and analyzed by FlowJo 10.8.1 (BD).

### Multiplex immunofluorescence staining

The sections undergo deparaffinization with BOND dewax solution and epitope retrieval process using a heat-induced method with BOND epitope retrieval solution or pH9 EDTA buffer. DAB staining for immunohistochemistry was performed to optimize antibody concentrations. The antibodies used in this analysis and the dilution factor were depicted in Supplementary table 2. Multiple cycles of heat-induced epitope retrieval, protein blocking, epitope labeling, and signal amplification were performed for multiplex staining, then the slides were counterstained using spectral DAPI, finally mounted using Proling Diamond Antifade Mountant (Thermo). Multispectral imaging was performed using the Vectra 3 Automated Quantitative Pathology Imaging System (Akoya). Firstly, whole slide images were acquired, and analyzed the tumor regions delineated by a certified neuropathologist at 20x of the original magnification. First, whole slide images were acquired after autoadjusting focus and signal intensity. Then, MSI was acquired in the tumor regions delineated by a certified neuropathologist at 20x of the original magnification. For analysis of MSI, we created a spectral library for all Opal dyes to subject acquired multispectral images to spectral unmixing that enabled the identification and separation of weakly expressing and overlapping signals from background to visualize the signal of each marker in inForm Tissue Finder software (inForm 2.6, Akoya Biosciences). Using InForm, the adaptive cell segmentation feature was used to identify the nucleus of the analyzed cells and to determine the nuclear and cytoplasmic compartments on each cell. A machine-learning algorithm within inForm was used in which cells were automatically assigned to a specific phenotype (GFAP+, TMEM119+, CD163+, CD16+, CD11c+, HLA-DR+). Batch analysis was used to analyze all tumor samples under the same segmentation and phenotype settings. The processing and analysis of images from all tumor samples were exported to cell segmentation tables. Exported files from inForm were processed in R using R packages Phenoptr and PhenoptrReports to merge and create consolidated single files for each tumor sample. Consolidated files had cell phenotypes as outputs that we employed for further quantification and spatial analyses using the Phenoptr R addin.

### RNA extraction and quantitative RT-PCR

Total RNA was extracted from cells by manual method using TRIzol (Invitrogen), and cDNA was synthesized from 1μg of total RNA using the LeGene 1st Strand cDNA Synthesis System (Legene Biosciences) according to the manufacturer’s instructions. Quantitative PCR was performed using TOPrealTM qPCR 2X PreMIX (Bio-Rad) and the CFX Connect Real-Time PCR Detection System (Bio-Rad). The primers used in this study are summarized in Supplementary Table 2.

### Western Blot

Cells were mixed with cell lysis buffer (Cell Signaling) containing protease and phosphatase inhibitor cocktail (Thermo Fisher Scientific) and heated for 10min in 95 °C. Cell lysates were resolved by sodium dodecyl sulfate-polyacrylamide gels, and separated proteins in gel were transferred to polyvinylidene difluoride membranes (Bio-Rad). Membrane was blotted with indicated antibody and immunoreactivity was detected with enhanced chemiluminescence solution (Thermo Fisher Scientific). Antibodies used in this study were summarized in Supplementary Table 3.

### Patient sample

The ethics approval was issued by the local committee of the University of Freiburg for data evaluation, imaging procedures and experimental design (Freiburg: protocol 100020/09 and 472/15_160880). All methods were carried out in accordance with the approved guidelines, with written informed consent obtained from all subjects. The patient provided preoperative (in the Department of Neurosurgery of the Medical Center - University of Freiburg, Freiburg im Breisgau, Germany) informed consent to take part in the study. Therapeutically resected access cortical tissue and glioblastoma tumor tissue were collected and transported to the laboratory immediately after surgical resection.

### Human organotypic slice culture system

Human organotypic slices were prepared and cultured as we previously described. Therapeutically resected access cortical tissue was immediately collected and immersed in the Preparation Medium saturated with carbogen (95% O2 and 5% CO2). Damaged tissue was dissected away from the primary tissue block. The primary tissue block was further dissected into smaller tissue blocks (6 x 6 mm^2^ dimension) for tissue sectioning. Sectioning was carried out using a Leica VT1200 semi-automatic vibratome (14912000001, Leica, Germany).

300 µm evenly thick cortical sections were collected in Growth Medium (GM) using a capillary glass pipette and settled on ice for 10 minutes for recovery. Recovered slices were plated on MilliporeTM inserts (PIPH03050, Merck KGaA, Germany) in 6-well plates (657160, Greiner BIO-ONE, Germany) supplied with GM and further cultured in the incubator at 37°C with 5% CO2 for following experiments. A number of 3 sections per culture well is recommended. The first medium refreshing was carried out 2 hours post tissue plating, followed by medium refreshing every 48 hours as a regular basis throughout the culture period.

### Autologous GB cell suspension

Patient-derived GB cell suspension was carried out, as we previously described using a Neural Tissue Dissociation Kit (T) (130-093-231, Miltenyi Biotec, Germany). Primary GB tissue obtained from the surgery was immediately processed and dissociated with the Neural Tissue Dissociation Kit (T) in C-Tubes (130-093-237, Miltenyi Biotec, Germany). Briefly, tissue was dissected into small pieces, incubated in enzyme 1 and enzyme 2 for 10 min with 2 min of tissue homogenizing in between each incubation. A filtering step followed to remove all visible tissue pieces with pia matter. Single cells in the filtered supernatant was further processed to remove myelin population along with red blood cells and non-viable cells. Eventually, viable single patient-derived GB cells in the final supernatant was seeded into T-75 cell culture flasks for cell culture or directly snap-frozen in cryopreservation medium in −80°C.

### GB cell labeling and detection assay

Cell TraceTM CFSE dye (C34570, ThermoFisher, USA) was used in this study to label the autologous GB cells. Tumor cells from cell culture were washed with PBS and detached with accutase for 5 min. CFSE dye was prepared with manufactor’s protocol. In a ratio of 1 µL dye per 2 x 10^6^ cells, the CFSE dye was used to label the tumor cells. Tumor cells were further observed under EVOS M7000 microscope (AMF7000, Thermofisher, USA) coupled with its onstage incubator system (AMC1000, ThermoFisher, USA) to confirm the success of fluorescent labeling for tumor cells.

### Personalized GB model

Personalized GB model system was generated based on the autograft of GB cells into the patient’s cortical sections. The CFSE labeled tumor cells were resuspended and inoculated into the patient’s cortical sections using a Hamilton micro syringe (80330, Hamilton, Switzerland) in the concentration of 20,000 cells per 1 µL.

### Tumor growth monitoring

Fluorescent images of sections after inoculation were acquired using EVOS M7000 microscope coupled with its onstage incubator system for monitoring tumor growth on a time-resolved manner of every 24 hours till the end point of the culture experiment.

### Fluorescence associated cell sorting

Fluorescence associated cell sorting (FACS) was performed to screen targeted cell/nucleus populations. In brief, tissue was homogenized and nuclei were isolated using Nuclei EZ Prep (NUC101-1KT, Merck KGaA, Germany) lysis buffer. Nuclei were labeled using DAPI (32670#5MG, Merck KGaA, Germany). Myelin and debris were removed by a sucrose gradient centrifugation step. 35,000 DAPI^+^ events per condition were collected from FACS sorter using 1.5 mL eppis coated with 100 µL 2% BSA.

### Single nucleus RNA-sequencing

Single nucleus RNA-sequencing, a droplet based approach, was performed according to the Chromium Next GEM Single Cell 3’ v3.1 protocol from 10X Genomics. Nuclei collected from FACS sorting were added to a prepared Master Mix and loaded in Chromium Controller for RNA recovery and generating GEMs. After reverse transcription and cDNA amplification, the enriched cDNA was fragmented, size-selected using SPRIselect (B23318, Beckman Coulter, USA), indexed using i7 index, and SI primer was added. The average length of final libraries was quantified using a 5200 Fragment Analyzer (M5310AA, Agilent, USA) with its HS NGS Fragment kit (DNF-474, Agilent, USA) and the concentration was determined using QubitTM 4 Fluorometer (Q33238, ThermoFisher, USA) with its 1x dsDNA HS kit (Q33231, ThermoFisher, USA). Diluted, pooled, and denatured final library was loaded in Illumina NextSeq 550 Sequencing System. NextSeq High Output kit v2.5 (20024906, Illumina, USA) was used in this study. Sequencing cycles for read1 – i7 – i5 – read2 were: 28 – 8 – 0 – 56.

### Spatially resolved transcriptomics

10 µm thick sections from fresh frozen tissue were mounted on specially designed spatially barcoded Visium Gene Expression Slide (1000188, 10X Genomics, Netherland). Mounted slide was fixed in 100% methanol following with H&E staining. EVOS M7000 microscope coupled with 20x magnificence lens was used to acquire bright field images. Post imaging processing was performed using FIJI ImageJ software. A permeabilization with pre-optimized incubation time was further performed to maximize capture oligo binding with sample mRNA. After reverse transcription, second strand cDNA was then cleaved off by denaturation. Following with a qPCR quantification using KAPA SYBR FAST qPCR Master Mix (KK4600, Roche, USA), cDNA was amplified and fragmented, and further size selected using SPRIselect. Similar to single nucleus RNA-sequencing library preparation, cDNA library quality was quantified using 5200 Fragment Analyzer., further indexed, amplified, and double-sided size selected following with an average base pair length quantification using 5200 Fragment Analyzer and a concentration quantification with Qubit^TM^ 4 Fluorometer. Final sequencing library was generated after a dilution, normalization and denaturation step. Illumina NextSeq 550 Sequencing System coupled with its NextSeq High Output kit v2.5 (20024904, Illumina, USA) was used in this study. Sequencing cycles for read1 – i7 – i5 – read2 were set up as 28 – 10 – 10 – 102.

### Statistical Analyses

Statistical analyses were performed using Prism Software 9.4.1 (GraphPad) and R Software 4.4.1 (R Studio). Unpaired Student’s t-test was used to compare statistical differences between the two groups. For Kaplan-Meier survival curves, the log-rank (Mantel-Cox) test was adapted to determine the significance between groups. Statistical significances were presented in P-value, or P < 0.05 was considered significant, *P < 0.05, **P < 0.01, ***P < 0.001, ****P < 0.0001.

## Discussion

Our study demonstrates the causal contribution of glioma-intrinsic MAPK/ERK signaling in modulating anti-tumoral immunity and responsiveness to immunotherapy. Our findings provide mechanistic insight into the previously observed association between MAPK-activating mutations, elevated p-ERK levels, and improved survival in GB patients treated with aPD-1 and aCTLA-4 immune checkpoint blockade therapy^9,10,21^. While MAPK signaling is known for its oncogenic and proliferative effects in cancer,^30^ our results reveal its crucial role in shaping the tumor-immune microenvironment, supporting the paradigm that molecular heterogeneity in GB tumors is linked to the variability in the immune microenvironment, TAM phenotypes, and ultimately, susceptibility to immunotherapy^11,41,42^.

Our study provides several lines of evidence that MAPK activation in GB cells enhances CD8^+^ T cell-mediated immune responses. Our CRISPR/Cas9 screen implicated the RAF-MEK-ERK branch of the MAPK pathway in glioma susceptibility to T cell recognition. This finding is corroborated by the differential responses to aPD-1 therapy in wildtype versus *Cd8* KO mice. The distinct phenotype of CD8^+^ T cells infiltrating the brains of long-term survivor mice with high p- ERK tumors treated with aPD-1 further supports the role of these cells in the observed anti-tumor immune response. Moreover, p-ERK promoted CD8^+^ T cell infiltration in mouse gliomas, with similar observations in human GB specimens. Consistent with this, clinical observations showed robust brain/lesional T cell infiltration in patients with elevated p-ERK who responded to aPD-1 therapy^9^. Additionally, transgenic murine glioma models, demonstrated that gliomagenesis in *Cd8* KO hosts, which lack CD8^+^ T cells, leads to tumors with elevated p-ERK, that are also more immunogenic^43^.

The mechanism by which MAPK/ERK signaling enhances T cell recognition of GB cells and response to immunotherapy appears to involve modulation of tumor cell immunogenicity and its relationship with the tumor-infiltrating microenvironment, particularly microglia. Our findings highlight the role of ERK phosphorylation in modulating both type I and type II IFN responses in glioma cells consistent with previous literature implicating MAPK/ERK signaling in the IFN response in other settings^31,44^. This regulation occurs through the activation of signaling pathways, including STAT1 and the MAPK/ERK pathway, which together drive the expression of genes involved in immune responses^45^. The mechanistic link between p-ERK and IFN responses in GB is supported by evidence showing that inhibition of ERK signaling impairs the induction of IFN response genes and dampens the overall immune response against tumors.

Furthermore, p-ERK was found to promote the expression of antigen presentation molecules in GB cells and surrounding microglia, likely related to the IFN response driven by MAPK/ERK signaling. This is consistent with antigen presentation being a downstream effect of IFN signaling^46–48^. The increase in antigen presentation, favorable microglia phenotype driven by MAPK/ERK signaling, as well as subsequent T cell infiltration and tumor recognition are consistent with mechanisms known to contribute to effective tumor cell killing^9^. The immune-modulating mechanisms driven by MAPK activation in GB cells translate into robust anti-tumoral immune responses beyond aPD-1 blockade, as evidenced by enhanced susceptibility to aCTLA-4 therapy. This suggests that MAPK/ERK signaling may serve as a predictor of response to various forms of GB immunotherapy and novel treatments that rely on T cell activity, such as vaccines, CAR-T cells, BiTEs^49–51^.

While MAPK activation contributes significantly to anti-tumoral immune responses, it appears to be necessary but not sufficient to elicit robust tumor rejection. Indeed, not all GB patients with elevated p-ERK benefit from aPD-1 therapy, whereas no patient with low p-ERK exhibited long survival after ICB^9^. Other factors, including the presence of tumor-specific antigens, bone marrow-derived immunosuppressive myeloid cells, T cell sequestration in the bone marrow, lymphopenia, and the use of steroids may also play critical roles, and influence tumor susceptibility to ICB. Additionally, several additional tumor-intrinsic features have been reported to modulate anti- tumoral immunity^10,18^.

While our study provides compelling evidence for the role of MAPK/ERK signaling in modulating GB immunogenicity and immunotherapy response, several limitations should be considered. Our experimental models, though informative, may not fully capture the complexity of human GB. Likewise, while our BRAF^V600E^ mutant GB slice culture model offers a unique platform to study tumor-microenvironment interactions, it lacks the systemic immune components and may not fully represent the *in vivo* tumor immune microenvironment. Additionally, while we observed a correlation between p-ERK levels and immunotherapy response in patients, the limited sample size necessitates validation through larger, prospective clinical studies across diverse patient populations. Despite these limitations, our multi-modal approach, combining *in vitro* studies, *in vivo* models, *ex vivo* human analyses, and large-scale omic approaches, provides a framework for understanding the role of MAPK/ERK signaling in GB immunobiology. Future studies addressing these limitations will be crucial for translating these findings into clinical applications and potentially paving the way for more effective, personalized immunotherapies in GB.

In conclusion, our study underscores the critical role of the RAF-MEK-ERK pathway in regulating immune responses in GB, offering new insights into the molecular mechanisms that drive immunotherapy responsiveness. By linking MAPK/ERK signaling to T cell-mediated tumor recognition and response, these findings offer valuable insights into the heterogeneity of GB and more importantly the potential of MAPK/ERK activation as a predictive biomarker and therapeutic target. While these findings offer promising avenues for personalized immunotherapy in GB, they also highlight the need for molecularly-guided approaches to address the heterogeneity of GB treatment responses, paving the way for more effective, tailored strategies in this challenging malignancy.

## Author contributions

K.S. Kim, C. Lee-Chang, D.H. Heiland and A.M. Sonabend conceptualized the project. K.S. Kim, Junyi Zhang, D.H. Heiland and A.M. Sonabend drafted the manuscript. K. Habashy, A. Gould, D. Chand, D. Levey, I. Balyasnikova performed the majority of the editing of the manuscript. The experiments presented were conducted and analyzed by K.S. Kim, K. Habashy, A. Gould, V.A. Arrieta, C. Dmello. L. Chen. J. Zhang, E. Grabis, J. Duffy, J. Zhao, Wenting Zhao, P. Canoll, P. A. Sims, R. Rabadan, and D. H. Heiland performed the bioinformatic analysis. V.A. Arrieta, A.M. Sonabend, S. Pandey, and Bin Zhang performed the histological analysis of the human tumors. C. Lee-Chang, D.H. Heiland, and A.M. Sonabend supervised the study.

## Acknowledgments and funding support

This work was supported by the NIH grants 1R01NS110703-01A1 (A.M.S., C.L.C.), 1U19CA264338-01 (A.M.S., C.L.C.), 1R01 NS122395 (I.V.B.), R37CA258426 (C.L.C), P50CA221747 SPORE for Translational Approaches to Brain Cancer (A.M.S., C.L.C., I.V.B.) and the Northwestern Nervous System Tumor Bank), as well as generous philanthropic support from the Sharon Moceri Foundation as well as Tina and Victor Kedaitis. R35CA253126 and Vagelos Precision Medicine Award (R.R. and J.Z.). Schematic illustrations are created with BioRender.com.

