## Supplementary figures for "MAPK/ERK signaling in gliomas modulates interferon responses, T cell recruitment, microglia phenotype, and immune checkpoint blockade efficacy"

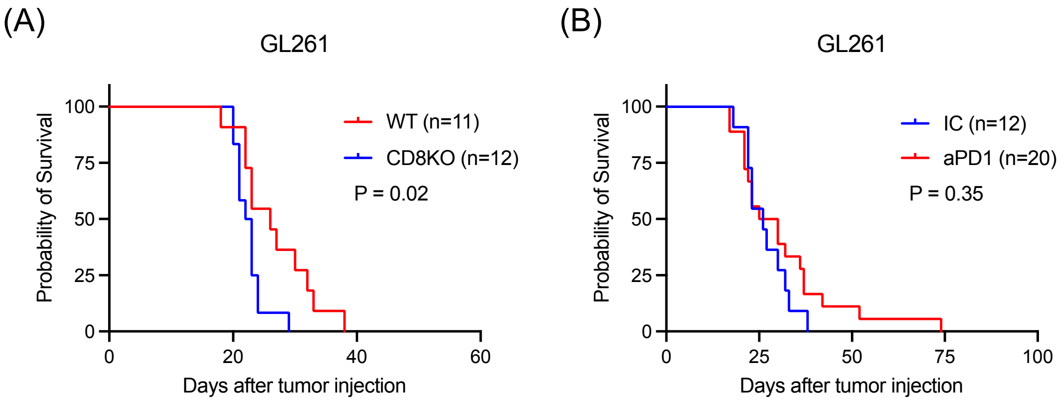


**Supplementary Figure 1.** (A) Kaplan-Meier survival curves from the kinome-wide CRISPR/Cas9 screen, comparing GL261-bearing C57BL/6 WT (n = 11) and *Cd8* KO mice (n = 12). (B) Kaplan-Meier survival curves from the kinome-wide CRISPR/Cas9 screen conducted in C57BL/6 WT mice comparing IC (n = 12) and aPD-1 (n = 20) treated groups. The P-values were calculated using the log-rank test.

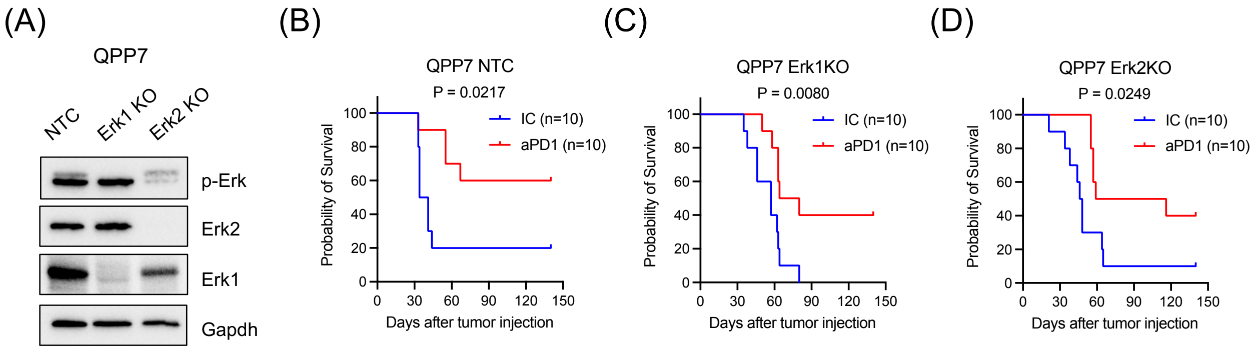


**Supplementary Figure 2.** (A) Western blot analysis showing KO of Erk1 or Erk2 in immunotherapy-sensitive QPP7 mouse glioma model. (B) Kaplan-Meier survival curves of QPP7-NTC bearing mice treated with IC (n = 10) or aPD-1 (n = 10). (C) Kaplan-Meier survival curves of QPP7-Erk1 KO bearing mice treated with IC (n = 10) or aPD-1 (n = 10). (D) Kaplan-Meier survival curves of QPP7-Erk2 KO bearing mice treated with IC (n = 10) or aPD-1 (n = 10). The P-values were calculated using the log-rank test.

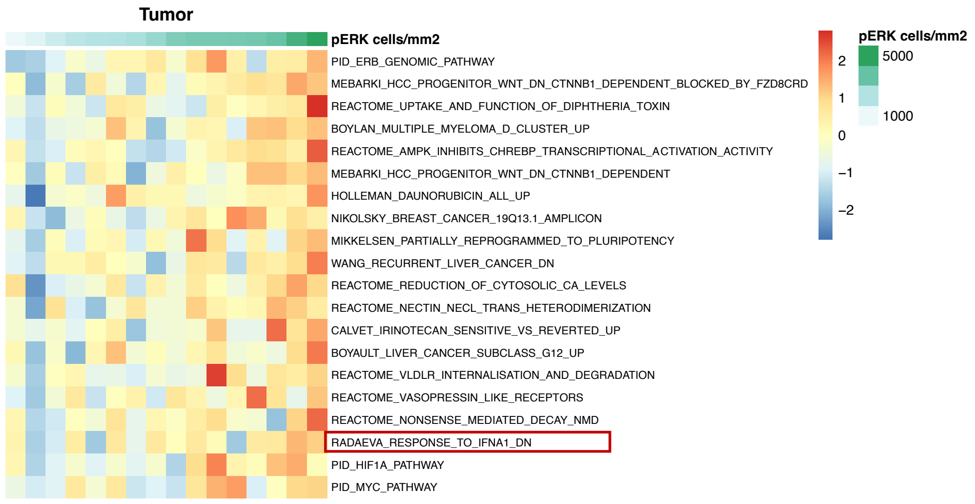


**Supplementary Figure 3.** Differential gene expression analysis of scRNA-seq data from human GB patient samples, categorized by continuous levels of p-ERK.

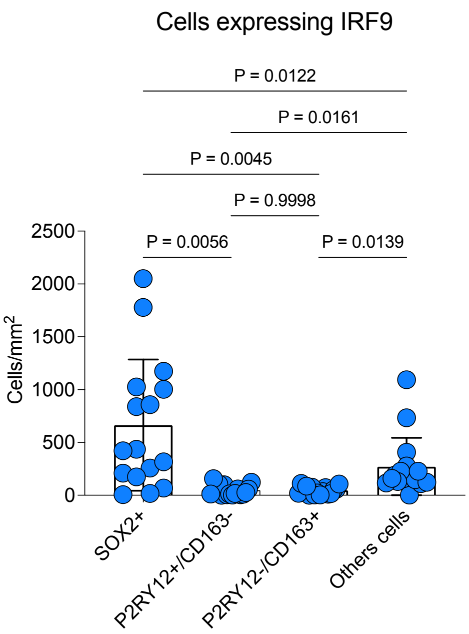


**Supplementary Figure 4.** Quantification analysis of immunofluorescence analysis staining for SOX2, P2RY12, CD163, IRF9, and DAPI in GB samples. One-way ANOVA was used to determine statistical significance, and corresponding P-values are shown in the figure.

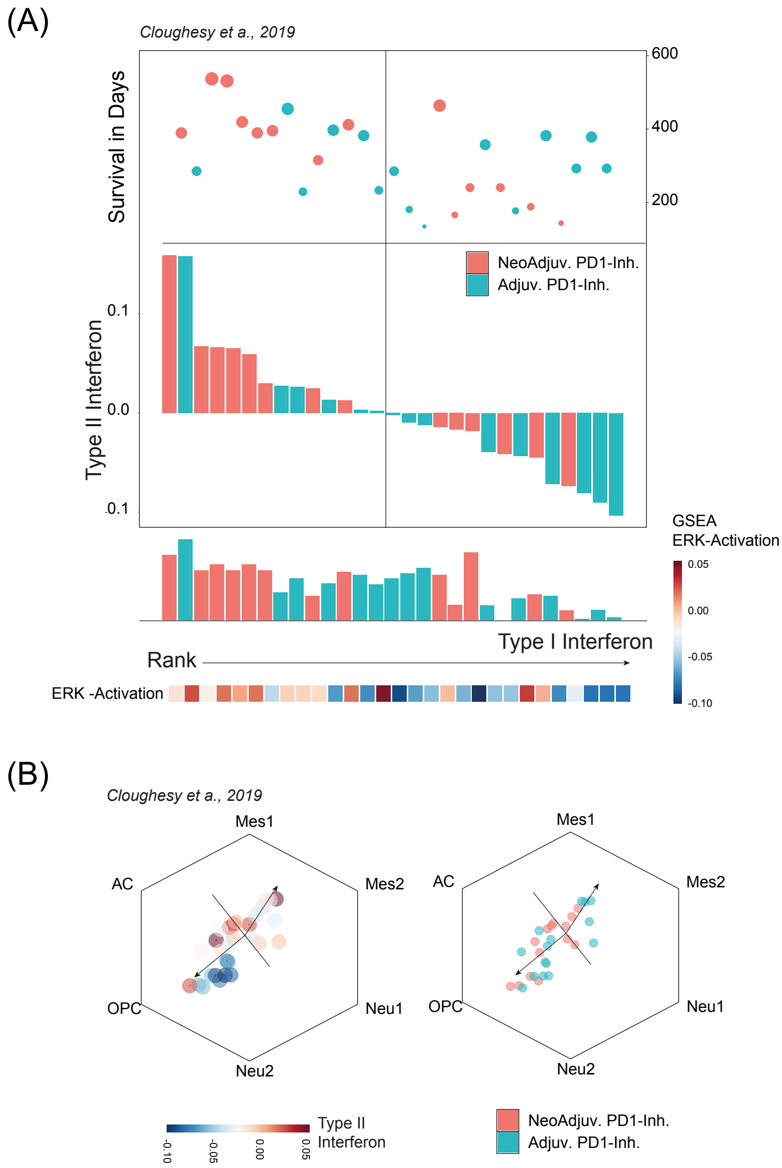


**Supplementary Figure 5.** (A) Analysis derived from Cloughsey's 2019 study, showing overall expression levels of Type I IFN across all cohorts and their respective survival rates. (B) The left panel depicts cell types and Type I interferon signaling, while the right side compares adjuvant PD-1 inhibitor therapy with neoadjuvant PD-1 inhibitor therapy.

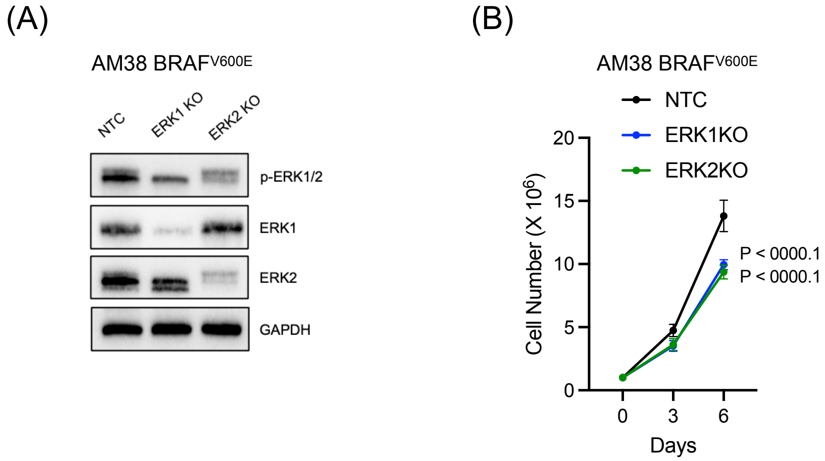


**Supplementary Figure 6.** Characterization of AM38-ERK KO cell lines. (A) Western blot analysis demonstrating the KO of ERK1 and ERK2 in the AM38 $BRAF^{V600E}$ cell line. (B) Cell proliferation assay showing the total cell number count for both ERK1 and ERK2 KO cell lines over time. The graph represents the mean ± SD (n = 4), one-way ANOVA was used to determine statistical significance, and corresponding P-values are shown in the figure.

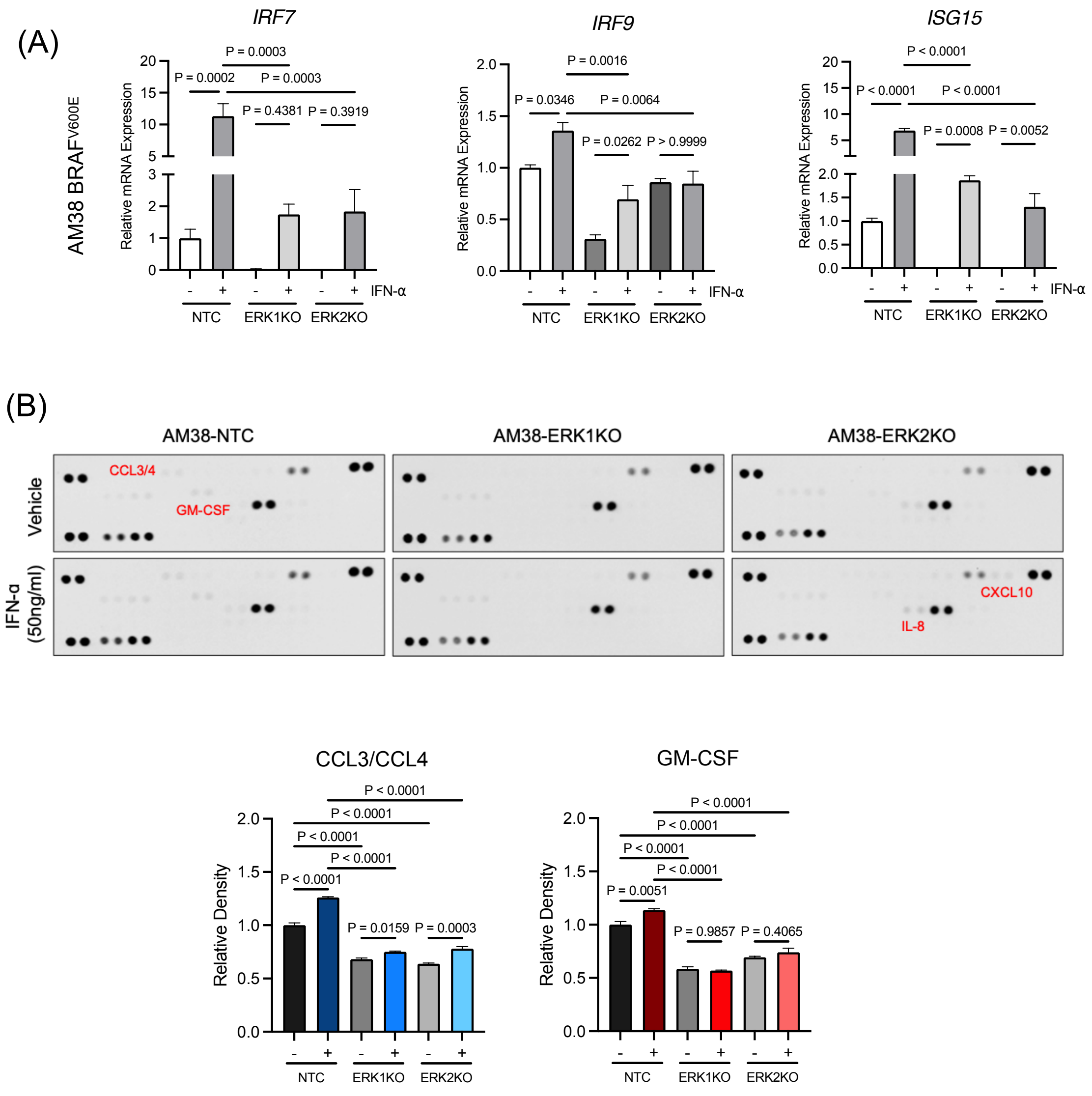


**Supplementary Figure 7.** (A) Quantitative RT-PCR results showing transcript levels of *IRF7*, *IRF9*, and *ISG15* in AM38-NTC, ERK1 KO, and ERK2 KO cells following exposure to IFN-α. Data are presented as mean ± SD (n = 4). (B) Cytokine secretion analysis using a human cytokine array kit (top panel). Bar graphs represent the relative fold change in CCL3/CCL4 (left) and GM-CSF (right) secretion from the culture supernatants of AM38-NTC, ERK1 KO, and ERK2 KO cells. Statistical significance was determined using one-way ANOVA, and P-values are indicated.

(A)

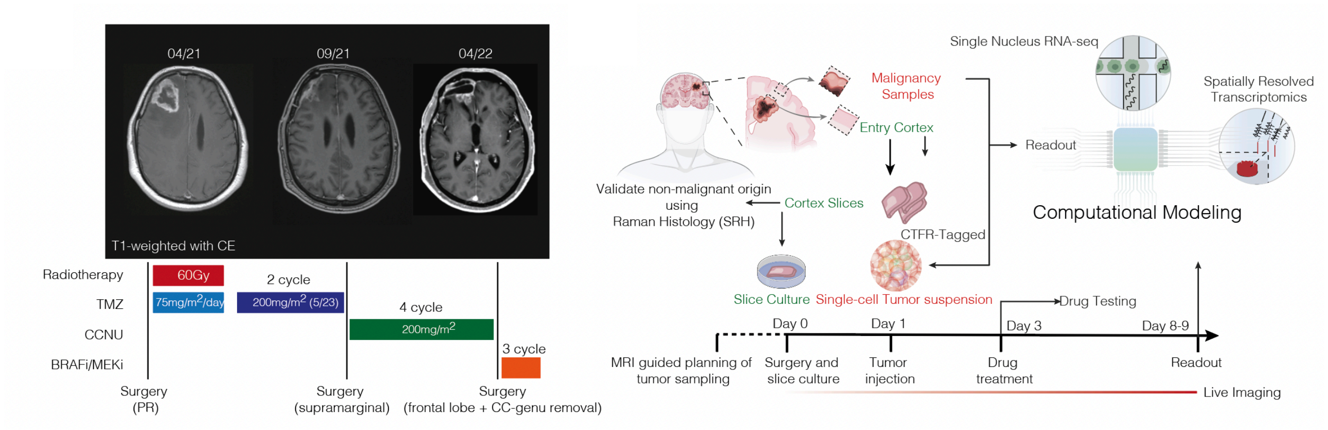


(B)

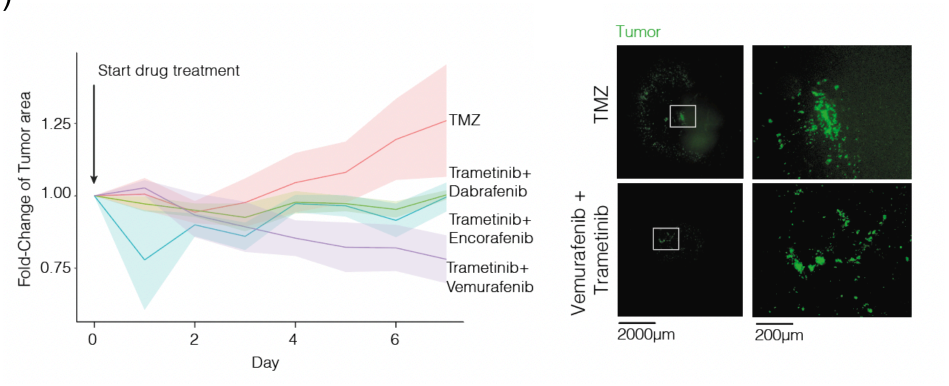


**Supplementary Figure 8.** (A) Brief illustration of autologous human neocortical slice model from BRAFV600E mutated GB patient. Tumor cells were implanted in cultivated cortex slices, then treated with temozolomide or BRAFi/MEKi combination.

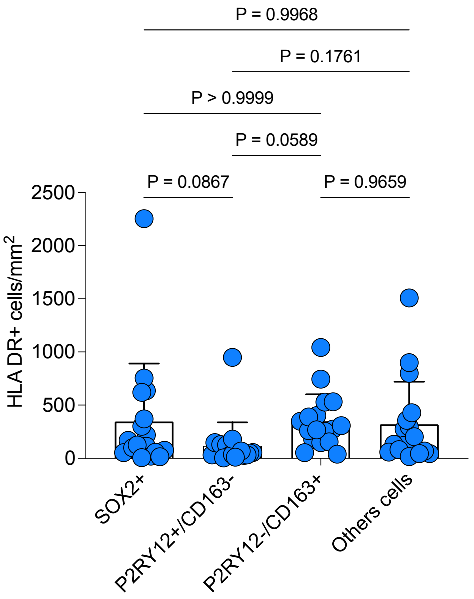


**Supplementary Figure 9.** Quantification analysis of immunofluorescence analysis staining for SOX2, P2RY12, CD163, HLA-DR, and DAPI in GB samples. One-way ANOVA was used to determine statistical significance, and corresponding P-values are shown in the figure.

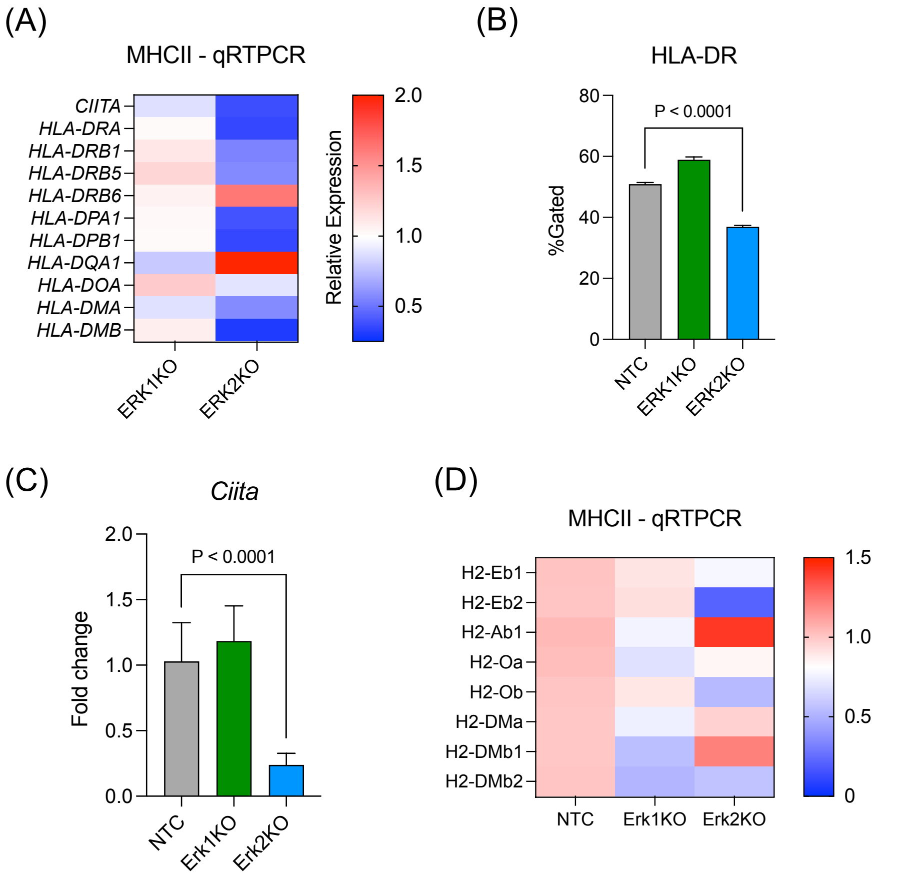


**Supplementary Figure 10.** (A) Quantitative RT-PCR results showing transcripts level of MHC class II genes comparing AM38-NTC cells and ERK1 KO or ERK2 KO cells. Data are presented as mean ± SD (n = 4). (B) HLA-DR protein expression was measured by flow cytometry-based analysis. Data are presented as mean ± SD (n = 4), unpaired two-tailed T test was used, and corresponding P-values are shown in the figure. (C) Quantitative RT-PCR results showing transcripts level of *Ciita* comparing QPP7-NTC cells and Erk1 KO or Erk2 KO cells. Data are presented as mean ± SD (n = 4), unpaired two-tailed T test was used. (D) Quantitative RT-PCR results showing transcripts level of MHC class II genes comparing QPP7-NTC cells and Erk1 KO or Erk2 KO cells. Data are presented as mean ± SD (n = 4), unpaired two-tailed T test was used, and corresponding P-values are shown in the figure.
